# Self-assembly of Grb2 meshworks revealed by Grb2-Gab1_497-528_ complex structure

**DOI:** 10.1101/2023.06.17.545433

**Authors:** Constanze Breithaupt, Tobias Gruber, Katharina Mandel, Marc Lewitzky, Annette Meister, Balbach Jochen, Stephan M. Feller, Milton T. Stubbs

## Abstract

The ubiquitously expressed adaptor protein Growth factor receptor bound protein 2 (Grb2) plays an essential role in signal transduction by binding to activated receptor tyrosine kinases through its SH2 domain and to downstream effectors via its N- and C-terminal SH3 domains (nSH3, cSH3). Here we present the first structure of ligand-bound full length Grb2. The crystal structure of Grb2 in complex with a bidentate nSH3-cSH3-binding peptide, derived from the multi-site docking protein Grb2- associated binder-1 (Gab1), provides molecular insight into effector recognition by Grb2 and reveals the assembly of a two-dimensional meshwork, consisting of multimeric filament-like Grb2 chains linked to each other by the bivalent bound Gab1_497-528_ peptide. Dominant contacts between Grb2 molecules in the multimer are provided by an intermolecular SH2/cSH3 domain interface that is also present in the closed dimer of ligand-free Grb2. We further show that Grb2 is able to self-assemble to form phase-separated condensates in solution. The Grb2 SH2 domain phosphotyrosine binding site is freely accessible in the multimeric assembly, and phase separation is fostered by addition of Gab1_497- 528_, as expected from the crystal structure. Multimeric assembly is also observed using a Grb2 SH2- cSH3 didomain construct, and suppressed using a Grb2 Tyr60Glu mutant, a mimic of the *in vivo* phosphorylated Tyr160 central to the SH2/cSH3 interface, demonstrating that an intact SH2/cSH3 interface is needed for Grb2 assembly in solution.

## Introduction

Growth factor receptor bound protein 2 (Grb2) is a ubiquitously expressed, highly conserved adaptor protein that transmits signals from activated receptor tyrosine kinases (RTKs) to downstream effectors. Discovered initially through its interaction with the tyrosine phosphorylated RTKs epidermal growth factor receptor (EGFR) and platelet-derived growth factor receptor (PDGFR)(1), Grb2 was subsequently found to be an essential component of the MAPK signaling cascade: bound to the guanine nucleotide exchange factor SOS, it translocates the latter to the plasma membrane upon EGFR activation, causing SOS-mediated activation of membrane-anchored Ras (2, 3). Numerous interaction partners of Grb2 have since been identified, including various RTKs, transmembrane and cytosolic adaptor proteins, GEFs, GTPases and E3 ubiquitin ligases (4-6) and Grb2 has been shown to participate in signaling processes that lead not only to cell growth and differentiation but also for example to actin cytoskeletal rearrangement and endocytosis (5, 6).

Grb2-associated binder-1 (Gab1), isolated from glial and medulloblastoma tumours, was identified as Grb2 binding protein [Holgado-Madruga et al., 1996]. Gab1 is the archetype for a family of proteins that play crucial roles in signal transduction, but whose aberrant expression or activation drives pathologies such as tumor development (7). A plethora of receptors (including c-Met, EGFR and gp130) and oncogenic fusion proteins (such as Bcr-Abl and Tpr-Met) rely on Gab proteins and their relatives for signal processing (8, 9), and a Gab1-Abl fusion protein was recently discovered in various tumors (10, 11). Gab1 consists of an N-terminal membrane binding Pleckstrin homology (PH) domain followed by a long intrinsically disordered region (IDR) of around 580 residues that contains docking sites for multiple signaling domains. It can thereby act as a dynamic assembly platform for large multi-protein complexes that integrate various intracellular signals transduced by diverse cell surface receptors (Fig. 1A) (9, 12-14).

**Figure 1.**
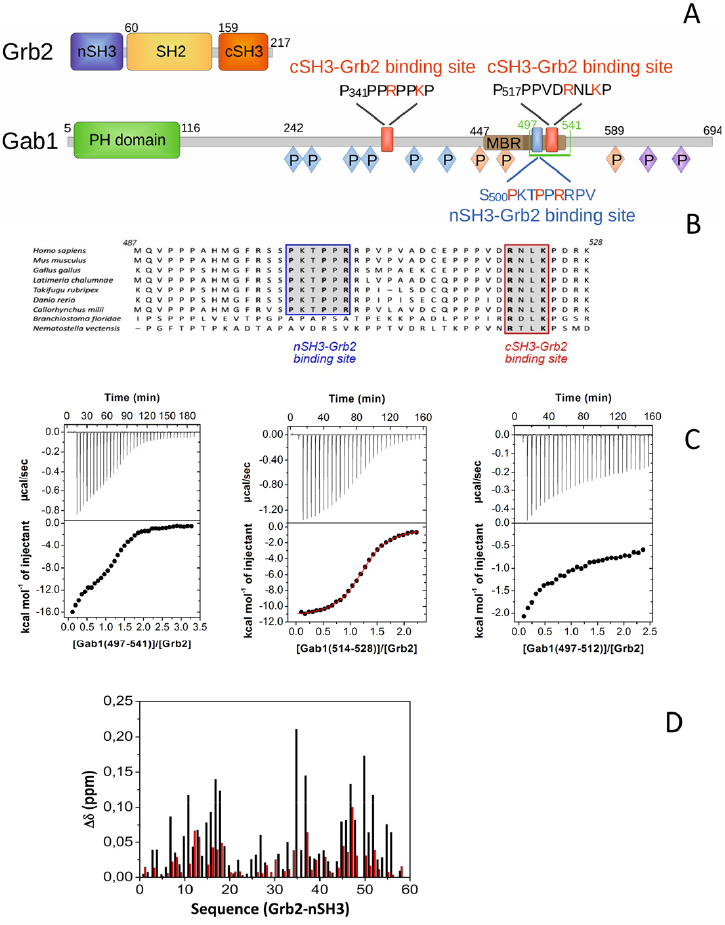
Grb2-Gab1 Interaction. **(A)** Schematic representation of structural features of human Grb2 and Gab1. P: tyrosine phosphorylation sites, bound by SH2-domains of Crk/CRKL (blue), PI3Kp85 (red), SHP2 (violet), MBR: c-Met binding region (brown), with sequence stretch essential for c-Met binding in dark brown. **(B)** Multiple sequence alignment of Gab1, shown for residues 487 – 528 (numbering according to human Gab1 sequence), including the second cSH3 Grb2 binding site and the preceding nSH3 Grb2 binding site, investigated in the present study. **(C)** ITC measurements of Gab1-peptides titrated to Grb2. Left: Gab1_497-541_ (containing both binding motifs), middle: Gab1_514-528_ (cSH3-Grb2 binding motif), right: Gab1_497-512_ (nSH3-Grb2 binding motif). **(D)** Chemical shift differences of ^1^H-^15^N-HSQC NMR-spectra of ^15^N-nSH3-Grb2 in the presence and absence of Gab1_497-541_ (red) and of the SOS variant peptide PPPLPPRRRR (black) (62).

Grb2 consists of an N-terminal SH3 domain (nSH3), a central SH2 and a second C-terminal SH3 domain (cSH3, Fig. 1A). The Grb2 SH2-domain binds to phosphorylated tyrosine residues embedded in the recognition sequence (pY)xNx (x being any amino acid) of RTKs or adaptor proteins. The two SH3-domains interact with downstream effector molecules; nSH3 prefers sequences that contain the classical PxxPxR class II consensus motif, whereas cSH3 binds preferentially to sequences that harbor the more recently discovered RxxK motif. The Gab1-Grb2 interaction was found to be dominated by binding of Gab1 to the cSH3 domain (15, 16). The interaction could be mapped to two regions in the long disordered tail of Gab1 that are conserved between Gab1 and Gab2 and bind to Grb2 cSH3 via their RxxK motif with dissociation constants in the low micromolar range (17, 18). In contrast, the proline-rich (PR) region of SOS1 contains four PxxPxR motifs that interact with the nSH3 domain of Grb2 in peptide binding studies (19).

Although the binding of short linear motifs to SH3 domains is well established, evidence has been accumulating that the specificity and affinity of many SH3-ligand interactions are dependent on factors outside of these. A number of structures of SH3 domains solved in complex with longer peptides show more extensive interactions, including regions outside the classical binding groove of the SH3 domain (20, 21). Mutagenesis and modelling studies indicate that extended binding modes also occur upon binding of Grb2 nSH3 to dynamin and of Grb2 cSH3 to a proline rich region at the C-terminus of FGFR (22-24). With its two SH3 domains, Grb2 can bind to hybrid peptide ligands created by joining nSH3- and cSH3-binding sequences via a suitable linker, with strongly elevated affinity compared to the monovalent interactions (25). Recently, three sites were identified in the PR region of SOS1 that interact with the cSH3 domain of Grb2 and that together with the known nSH3 binding regions of SOS1 allow several different bivalent binding modes between SOS1 and Grb2 (26). This bivalent binding was suggested to contribute to the more than 100 fold increase in affinity observed for the interaction of full length SOS1 with full length Grb2 compared to SOS1 peptide – Grb2 nSH3 interactions (26-28).

The 3.1 Å crystal structure of full length ligand-free Grb2 shows a compact dimeric arrangement of two Grb2 molecules that is in line with Grb2 forming a weakly associated dimer in solution (25, 29-31). Upon addition of SH2-binding phosphotyrosine ligands, the Grb2 dimer dissociates into rather extended monomers exhibiting variable domain orientations (25, 30). Grb2 Tyr160, positioned within the dimer interface, is phosphorylated *in vivo* by RTKs like FGFR2 or c-Met and the presence of phosphorylated Tyr160 or its mimic glutamate in the Tyr_160_Glu Grb2 mutant inhibits Grb2 dimerization (30, 32, 33). The fact that several oncogenic fusion tyrosine kinases such as Bcr-Abl, NPM-ALK and TEL-JAK2 also phosphorylate Tyr160, that high levels of phosphorylated Tyr160 have been detected in high grade pre-metastatic tumors but not in less malignant tumors or normal tissue, and that the degree of Tyr160 phosphorylation correlates with the progression of colon and prostate cancer, suggest a role of oligomeric Grb2 in cancer progression (30, 33).

Many signaling cascades rely on the formation of supramolecular intracellular assemblies that allow highly specific binding and spatiotemporal regulation of signaling (34, 35). Recent studies indicate that signaling complexes from a number of pathways compartmentalize into protein-enriched condensates formed by liquid liquid phase separation (LLPS) (35). One such example is the two-dimensional network formed at the membrane, early in T-cell signaling by multivalent interactions of Grb2, SOS1 and the transmembrane adapter LAT, which exhibits the liquid-like dynamic properties of phase separated biological condensates (28, 35, 36). Multimerization is initiated by phosphorylation of the intrinsically disordered region of LAT, induced by T-cell receptor activation, and subsequent interaction of LAT with Grb2 via three Grb2-specific phosphotyrosine binding motifs. Both Grb2 SH3 domains and at least two Grb2 binding motifs on SOS1 were shown to be required for cluster formation (28, 37).

Grb2 plays a central role in this process, as phase separated clusters can be formed from Grb2 and phosphorylated LAT alone or from Grb2 and the phosphorylated cytoplasmic tail of EGFR (38). Phosphorylation of Grb2 Tyr160 or mutation to glutamate inhibited phase separation, suggesting that the Grb2 dimer interface is also needed for EGFR:Grb2 clustering. Activation of Ras as well as phosphorylation of the downstream kinase ERK were shown to depend on the degree of EGFR- and of LAT-clustering, respectively, underlining the physiological relevance of the observed multimerization (37, 38).

Our understanding of Grb2 structure, limited currently to those of ligand-free full-length Grb2 and of isolated Grb2 domains in presence or absence of short peptide ligands, lags behind the impressive recent biochemical data on the complex interactions experienced by Grb2 (24, 26, 28, 35-38) and fails to explain the observed multivalency. Here we present the identification of a nSH3-Grb2 binding region on Gab1 and the crystal structure of full-length Grb2 in complex with the bidentate nSH3-cSH3-binding peptide Gab1_497-528_. The structure reveals the assembly of a two-dimensional network consisting of multimeric Grb2 filament-like chains linked to each other by the bivalent bound Gab1_497-528_ peptide. We show that Grb2 is able to self-assemble and to form phase-separated condensates in solution, a process that is fostered by addition of Gab1_497-528_ and an assembly that is compatible with phosphotyrosine ligand SH2 binding.

## Results

### Identification of an nSH3-Grb2 binding site in Gab1

When analyzing sequence variations of Gab1 proteins, we observed that the second of the two Grb2-cSH3 binding regions of Gab1 (P_516_PPVD**R**NL**K**P_525_) is preceded by the class II consensus motif **P**_501_KT**P**P**R**_506_ that is not present in other members of the Gab family but has been conserved in Gab1 for a period of ∼ 500 million years of evolution (Fig. 1B). We synthetized Gab1 peptides containing one (Gab1_497-512_, Gab1_514-528_) or both (Gab1_497-541_) binding motifs and analyzed their interaction with Grb2 using ITC. The ITC curve for Gab1_497-541_ binding to Grb2 shows two separated binding events, suggesting that both motifs participate in the interaction (Fig. 1C). Titration with a Gab1 peptide in which the PxxPxR motif has been mutated, or with the shorter peptide Gab1_514-528_ that contains only the RxxK motif, resulted in a single-site binding curve, confirming that the N-terminal PxxPxR motif is involved in binding of Gab1_497-541_ to Grb2 (Fig. 1C, S1). The observed dissociation constant of 4 μM for the C-terminal Gab1_514-528_ peptide is in agreement with the *K*_D_ value reported for the interaction between the homologous Gab2 peptide and the isolated Grb2-cSH3 domain (18). Titration of the long peptide Gab1_497-541_ to Grb2-cSH3 did not alter the binding curve significantly, indicating that the cSH3 domain binds the RxxK motif alone and not the PxxPxR motif (Fig. S1). Titrating the short N-terminal peptide Gab1_497-512_ which only contains the PxxPxR binding motif to Grb2 resulted in a complex non-sigmoidal curve, indicating that Gab1_497-512_ binds to the nSH3 domain of Grb2 but that additional processes accompany binding (Fig. 1C). ^1^H-^15^N-HSQC NMR-spectra of the isotope labeled Grb2-nSH3 domain were analyzed in the presence and absence of Gab1_497-541_ and compared with corresponding spectra in the presence of a Grb2-nSH3 binding SOS-peptide (Fig. 1D). The chemical shift differences observed after addition of Gab1_497-541_ showed that essentially the same Grb2-nSH3 residues are involved in the binding of both peptides.

The nSH3 ^1^H-^15^N-HSQC NMR-spectra exhibited an additional set of poorly dispersed signals compatible with a fraction of nSH3 molecules in a disordered state. This is in line with the reported low stability of nSH3 of the *Drosophila* Grb2-homolgue Drk, for which a 30% unfolded fraction in equilibrium with the folded state was observed by NMR, with the equilibrium shifting upon binding to the HtrA2 protease (39). Such a coupling of folding and binding might explain the complex ITC binding curve of Grb2-nSH3 to Gab_497-512_ (Fig. 1C).

### Crystal structure of the Grb2-Gab1_497-528_ complex

We tested a number of Gab1 peptides containing both SH3 binding motifs for their crystallization behavior in complex with Grb2, and obtained crystals in the presence of Gab1_497-528_. The Grb2-Gab1_497-528_ complex crystallized in space group P2_1_ with one complex per asymmetric unit. Diffraction data could be collected to a limiting resolution of 2.2 Å and the structure was refined to R/R_free_ values of 19.9%/22.6% (Table 1). The polypeptide chain of Grb2 is clearly defined except for residues 153-156 in the linker region between the SH2 and cSH3 domains, and five C-terminal residues. For Gab1_497-528_, no electron density could be attributed to four terminal residues, nor to residues 512-514 in the linker between the two SH3-binding regions. The three individual Grb2 domains show high structural similarity to those in ligand-free full-length Grb2 (29) (rmsd of superimposed atoms: 0,49 Å – 0,62Å), but the relative domain arrangement is significantly different: Grb2 adopts an extended open conformation, and the Grb2 dimer seen for the apo structure is not present in the Grb2-Gab1_497-528_ complex crystals. Interestingly, Gab1_497-528_ bridges two Grb2 molecules by binding to the Grb2-nSH3 domain via residues S_500_PKTPPRRPV_509_ and to the cSH3 domain of a neighboring Grb2 molecule with residues P_516_PPVDRNLKP_525_ (Fig. 2A-C).

**Figure 2.**
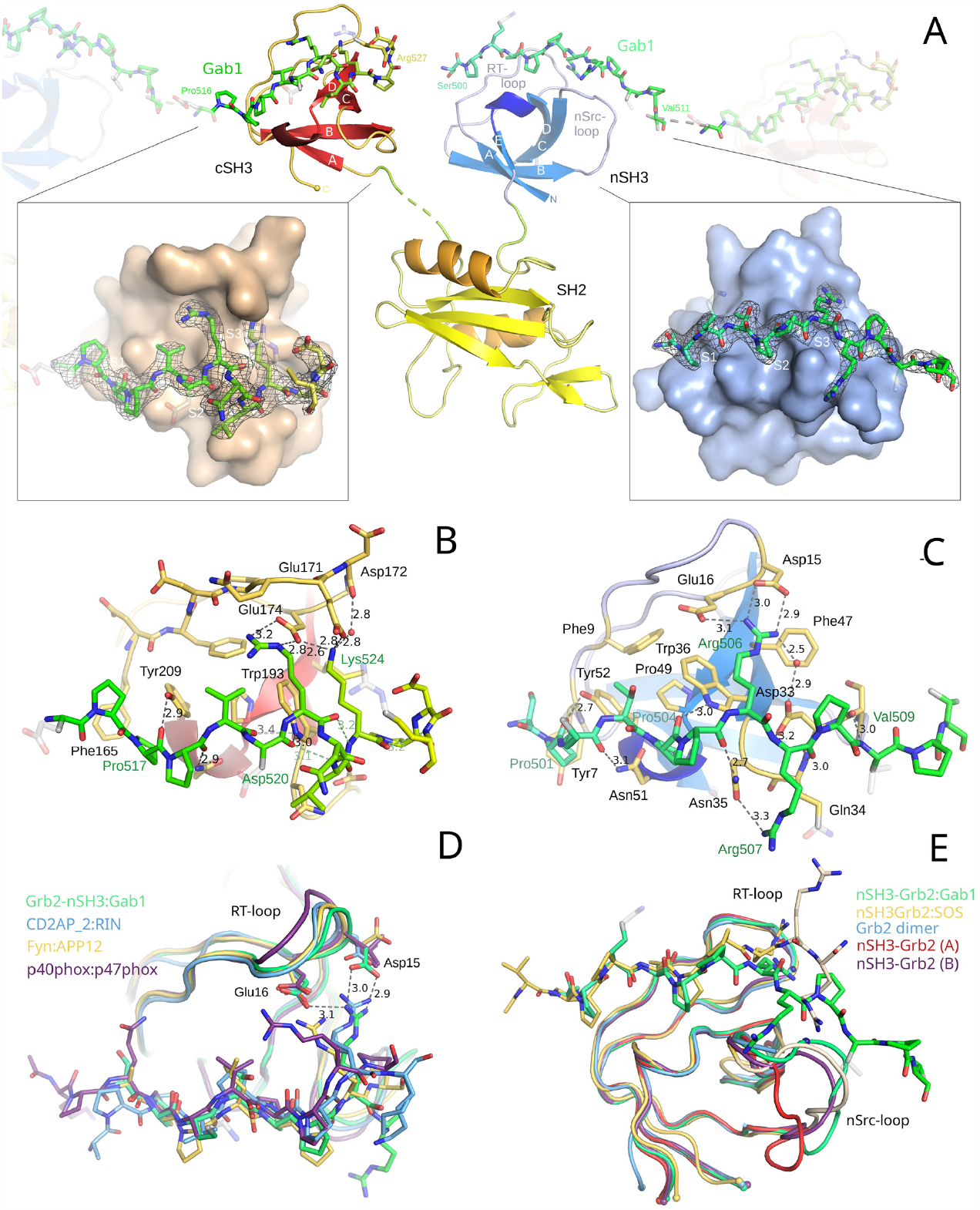
Three-dimensional structure of the Grb2-Gab1_497-528_ complex. The secondary structure elements and the surfaces of nSH3- and cSH3-Grb2 are depicted in blue and red, respectively. Gab_497-528_ is shown as stick model with a color gradient from N- to C-terminus from turquoise to chartreuse. Atoms that are not visible in the electron density are colored white. **(A)** Overall structure of one Grb2 molecule in complex with two Gab1_497-528_ peptides. Peptides residues 512-515 are not ordered in the structure. The molecular surface of cSH3- and nSH3-Grb2 and the 2F_o_-F_c_ electron densities of the bound peptide residues are shown in the left and right boxes, respectively. The cavities of the S1, S2 and S3 sites are marked. **(B)** Ligand binding site of cSH3-Grb2 with bound Gab_497-528_. Hydrogen bond lengths are given in Å. **(C)** Ligand binding site of nSH3-Grb2 with bound Gab_497-528_. **(D)** Structural superposition of the Cα-traces of nSH3-Grb2 with the ligand-bound SH3 domains of CD2AP (second SH3 domain), Fyn and p40phox (50, 63, 64). For clarity, the non-binding terminal ligand residues are not shown. The strong hydrogen bonds of nSH3-Grb2 Arg506 to Asp13 and Glu16 are depicted with lengths given in Å. The comparison shows a highly conserved mode of ligand binding to the S1 and S2 subsites and more diverse interactions in the S3 subsite. Similar to Gab1 Arg506, the corresponding Arginine of Rab Interactor 3 (RIN3) binds to the respective aspartate and glutamate residues at the tip or the RT-loop of CD2AP-2, whereas the corresponding arginines of APP12 and p47phox only interact with a single acidic residue of FynSH3 and p40phox-SH3, respectively. NMR-studies of the SOS1 V_1149_PPPVPPRRR_1158_ peptide in complex with Grb2 nSH3 have shown S1/S2 – peptide interactions that closely resemble the observed Gab1 binding, participation of Arg1156 (Arg506 in Grb2) in fluctuating hydrogen bonds and high flexibility of the two terminal arginines (see also **(E)**) (47). The observed Gab1 S_500_PKTPPRRPV_509_ – Grb2 nSH3 binding mode suggests that the hydrogen bonding partners of the SOS1 Arg1156 side chain are Asp15 and Glu16 of Grb2, in line with NMR data and recent modelling studies (22, 26, 49). **(E)** Structural superposition of the Cα-traces of nSH3-Grb2 in complex with Gab_497-528_, the minimized average NMR structure of the nSH3-Grb2-SOS1_1149-1158_ complex (yellow) and the crystal structures of the nSH3 domain of ligand-free Grb2 (blue) and of the two molecules in the asymmetric unit of ligand-free nSH3-Grb2 (red, chain A and violet, chain B) (29, 46, 47). In the ligand-free nSH3-Grb2 crystal structure, the proline-containing C-termini of two neighboring nSH3 domains interact with the binding site of chain B nSH3 by mimicking S1/S2-binding and S3/nSrc-loop-binding, respectively. Residues of the nSH3-Grb2-SOS1_1149-1158_ complex that show strong variation within the NMR structure ensemble are depicted in beige. The Cα-atoms of Grb2 Asp15 in the RT- and Asp33 in the nSrc-loop are shown as blue spheres. The movement of the nSrc-loop, observed in molecular dynamics simulations of Grb2-nSH3 upon complex formation with nSH3 binding SOS1 peptides, led to a reduced Asp15(Cα) - Asp33(Cα) distance. In the Gab1-Gbr2 structure, the nSrc-loop of nSH3 is positioned near the RT-loop with a small Asp15(Cα)-Asp33(Cα) distance of 12.4 Å, supporting the results of the molecular dynamics calculations.

This first crystal structure of Grb2-nSH3 in complex with a ligand provides valuable insights into its effector recognition. The nSH3 domain consists of two antiparallel three-stranded β-sheets (E-A-B_shared_ and B_shared_-C-D), that pack against each other at an angle of approximately 90°, creating the sandwich structure characteristic for SH3 domains. The ligand-binding groove is formed by residues of strands C and D, of the adjacent nSrc-loop and 3_10_-helix, and of the long hairpin-like like RT-loop (Fig. 2A). Within the groove, residues Tyr7, Asn51/Tyr52, and Asn35/Trp36 are arranged like rungs of a ladder to create three subsites for the consensus motif PxxPxR (Fig. 2A,C): the hydrophobic shallow binding pockets S1 and S2 that each bind preferentially to xP-dipeptides, and the “specificity region” S3 that binds the basic Arg/Lys. In accordance with other type II consensus peptide-SH3 crystal structures (Fig. 2D), Gab1_497-528_ peptide residues S_500_PKTPP_505_ adopt a polyproline II (PPII) helical conformation, with Ser500/Pro501 and Thr503/Pro504 occupying the S1 and S2 sites of Grb2, respectively. The four main chain carbonyl-oxygen atoms of the PPII-helix that point towards the binding groove are all involved in hydrogen bonds to the side chains of the “ladder residues” Asn51, Tyr52, Asn35 and Trp36. Pro501 and Pro504 make hydrophobic interactions within the binding pockets, as do the aliphatic portions of Ser500 and Thr503 whose hydroxy groups point towards the solvent. Basic residue Arg506 binds to the S3 site by stacking onto Trp36 and by forming three hydrogen bonds between its guanidyl moiety and the carboxylate groups of Asp15 and Glu16 in the Grb2 nSH3 RT-loop, as well as to nSrc-loop Asp33 via a bridging water molecule. Interestingly, the interaction of Gab1_497-528_ with the Grb2 nSH3 domain extends beyond the consensus motif through contacts between Gab1 residues R_507_PV_509_ and nSrc-loop residues Ser32-Asn35, with main chain hydrogen bonds between Arg507-Gln34 and Val509-Ser32. This latter interaction is stabilized through stacking of Arg507 against Gln34 and a weak hydrogen bond between its guanidinium group and Asn35.

The interaction of the C-terminal Grb2 cSH3 domain with Gab1 residues P_516_PPVDRNLKP_525_ follows the binding mode of RxxK-motif ligands and is very similar to the binding of Gab2_510-519_ to Grb2-cSH3 (18), with Asp520 (Asn in Gab2) being the only difference in sequence in this region (Fig. 2B). Whereas Gab1 Pro517/Pro518 bind to the cSH3 S1 site as typical xP dipeptides, the S2 site is only partly occupied with Val519. Asp520 deviates from the cavity, with its disordered side chain oriented towards the solvent, bringing Arg521 in the correct position to bind to the S3 site via two hydrogen bonds to Glu174 of the RT-loop. As seen for Gab2_510-519_, Gab1 residues R_521_NL_523_ form a 3_10_-helix so that Lys524 is oriented parallel to Arg521 and able to form two strong hydrogen bonds to the side chains of Grb2 Glu171 and Glu174 as well as a water-mediated hydrogen bond to the main chain of Asp172 in the RT-loop (Fig. 2B).

### Filament-like multimerization of Grb2 in crystal structure

Inspection of inter-protein contacts within the crystal reveals that Grb2 forms filament-like multimers oriented along the crystallographic twofold screw axes (Fig. 3A). The Gab1_497-528_ fragment is not involved in intra-multimer contacts but bridges two Grb2-multimers by binding with one SH3-binding motif to one multimer and with the second to a neighboring Grb2-multimer. 1890 Å^2^ of the solvent accessible surface are shielded by one Grb2-Grb2 interaction in the multimer, which is in the range observed for protein-protein complexes (40). Small interfaces that lack intense interactions involving only 15 amino acids in total are found between the C-terminal end of the SH2-domain and the nSH3*-as well as cSH3*-domain of the protomer (Grb2*) adjacent to the Grb2 protomer in the multimer chain (Fig. 3B). The most prominent interface is found between the AB-β-sheet and the CD-loop of the cSH3 domain with the complete N-terminal face of the SH2*-domain of a neighboring Grb2 molecule, including SH2*-residues preceding α-helix αA*, the AB*-loop, the small DE*-β-sheet and residues in the N-terminal region of α-helix αB*. cSH3-Phe182 binds extensively to a depression in a highly hydrophobic patch of SH2, formed by residues Trp60*, Phe95*, Ile128* and by Phe61* that π-π-stacks with Phe182. The side chain of cSH3-Gln162, fixed by a strong hydrogen bond to SH2*-Asn129* and an intramolecular hydrogen bond, stacks to the other side of Phe182 to result in nearly complete burial of the aromatic surface residue. At the edge of the hydrophobic interface, cSH3-Tyr160 binds to SH2*-Phe61* and makes weak polar contacts to Glu87* and Lys64* via its hydroxy group. These interactions are supplemented by an adjoining polar region in SH2*, governed by eight hydrogen bonds from cSH3 residues Asp181, Arg179 and Arg178 to SH2* residues Tyr118*, Arg112*, Gly116* and Asn126*. Comparing the present structure with that of ligand-free Grb2, which crystallized as a compact dimer with two Grb2 molecules related by a twofold symmetry axis, it is clear that the intermolecular interface involves exactly the same SH2 / cSH3* domain contacts in the Grb2 dimer as in the Grb2 multimer structure. Further interactions that support the observed dimerization are primarily formed between the two cSH3 domains, the linker regions and the two C-termini.

**Figure 3.**
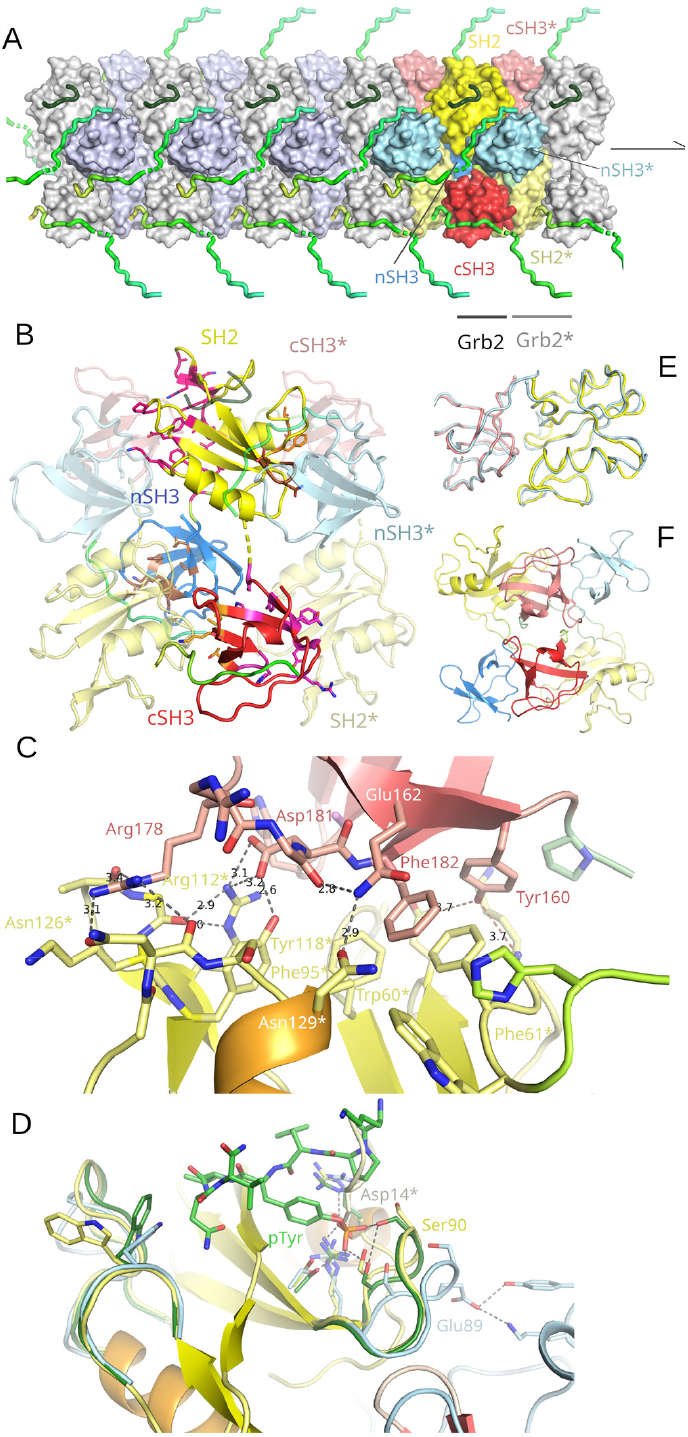
Three-dimensional structure of multimeric Grb2. **(A)** The filament-like Grb2 structure shown as surface representation. One Grb2 molecule is colored according to its domains and the two adjacent Grb2 molecules are depicted in lighter colors. The Gab peptides are shown as green ribbon structures. The position of the SH2-Grb2 binding sites is visualized by the ribbon structure of the Bcr-Abl_174-180_ peptide (dark green) obtained by structural superposition of the SH2-Grb2 Bcr-Abl_174-180_ complex structure onto the SH2 domain of the Grb2-Gab1_497-528_ complex (42). **(B)** Cartoon representation of one Grb2-Gab1_497-528_ complex and two neighboring complexes. Residues that contribute to the large SH2-cSH3 interface are shown as pink sticks, residues of the small interface in brown (SH2-nSH3) and orange (cSH3-SH2, “backward” interaction). **(C)** Close-up view of the SH2-cSH3 Grb2 multimer interface. Lengths of hydrogen bonds between residues of the SH2 (yellow) and cSH3 (red) domain are given in Å. **(D)** Structural superposition of the SH2 domains of Grb2-Gab1_497-528_ (yellow), ligand free Grb2 (dimer, light blue) and SH2-Grb2 in complex with Bcr-Abl_174-180_. Asp14 of a neighboring Grb2 molecule that mimics the ligand’s phosphotyrosine in the Grb2-Gab1_497-528_ crystal is shown as white stick model. **(E)** Structural superposition of the Cα-traces of the intermolecular SH2 (yellow) - cSH3* (red) domain complex of Grb2-Gab1_497-528_ with those of the interacting SH2 and cSH3* domains of the two Grb2 protomers in the Grb2 dimer (light blue)(29). **(F)** Cartoon representation of the Grb2 dimer viewed along the twofold symmetry axis. The SH2-cSH3* interaction corresponds to the interaction in the Grb2 multimer whereas all other interactions differ.

The SH2 peptide-binding site of Grb2, located in the monomer at the tip of the central β-sheet and delimited by the BC-loop on one side and by the specificity determining EF loop on the other, is solvent exposed and freely accessible in the Grb2 multimer (Fig.1A,3A). Ligands that bind to the Grb2 SH2 domain do not bind in the extended conformation typical of SH2 binders but adopt a type I β-turn conformation that positions the asparagine side chain of the (pY)xNx consensus motif in a binding pocket close to the phosphotyrosine binding site (41, 42). The SH2-binding site does not appear to be affected by Grb2 multimerization, and exhibits a backbone conformation that is highly similar to that of ligand bound Grb2 SH2-domains. Asp14* of a neighboring Grb2 filament protomer, part of a minor crystal contact, mimics phospho-tyrosine binding by inserting deeply into the phosphate binding pocket to form hydrogen bonds to the SH2-residues Arg67, Arg86 and Ser88, indicating that Grb2 SH2 ligands can bind to multimeric Grb2 (Fig. 3D). In contrast, the BC-loop curves away from the phosphotyrosine binding site in the Grb2 dimer structure, which allows Glu89 of the BC-loop to form hydrogen bonds to Tyr7 and Lys6 of the dimeric counterpart, but simultaneously disrupts the phosphate binding pocket by impeding the formation of three strong hydrogen bonds between the tyrosine phosphate group and Ser88/Ser90. This structural difference could contribute to the Grb2 dimer dissociation observed after addition of Grb2-binding phosphotyrosine ligands.

### Self-assembly of Grb2 in solution mediated by SH2/cSH3 interactions

When handling concentrated solutions of Grb2, we had noted that these turned turbid upon cooling. We therefore performed temperature-dependent DLS measurements on concentrated solutions of Grb2. At 25°C, Grb2 exhibited a mean hydrodynamic radius of 3.4 nm, which is in the range expected for monomeric/dimeric Grb2. At 15°C, however, a high molecular weight species with a mean hydrodynamic radius of 1.2μm appeared that became the only species at lower temperatures (Fig. 4A). Increasing the temperature again reverted the process completely, demonstrating that Grb2 reversibly self-assembles at high concentrations in an enthalpy-driven process. Using polarized light microscopy, we observed that Grb2 phase-separated during cooling to form μm-sized droplets, in line with the particle radii observed in DLS (Figure 4B). Thus, Grb2 can undergo homotypic LLPS without the participation of other proteins. Turbidity measurements at different Grb2 concentrations showed that the cloud point temperature at which LLPS begins decreases upon lowering the Grb2 concentration, as expected for systems that phase separate upon cooling (Fig. 4C) (43-45).

**Figure 4.**
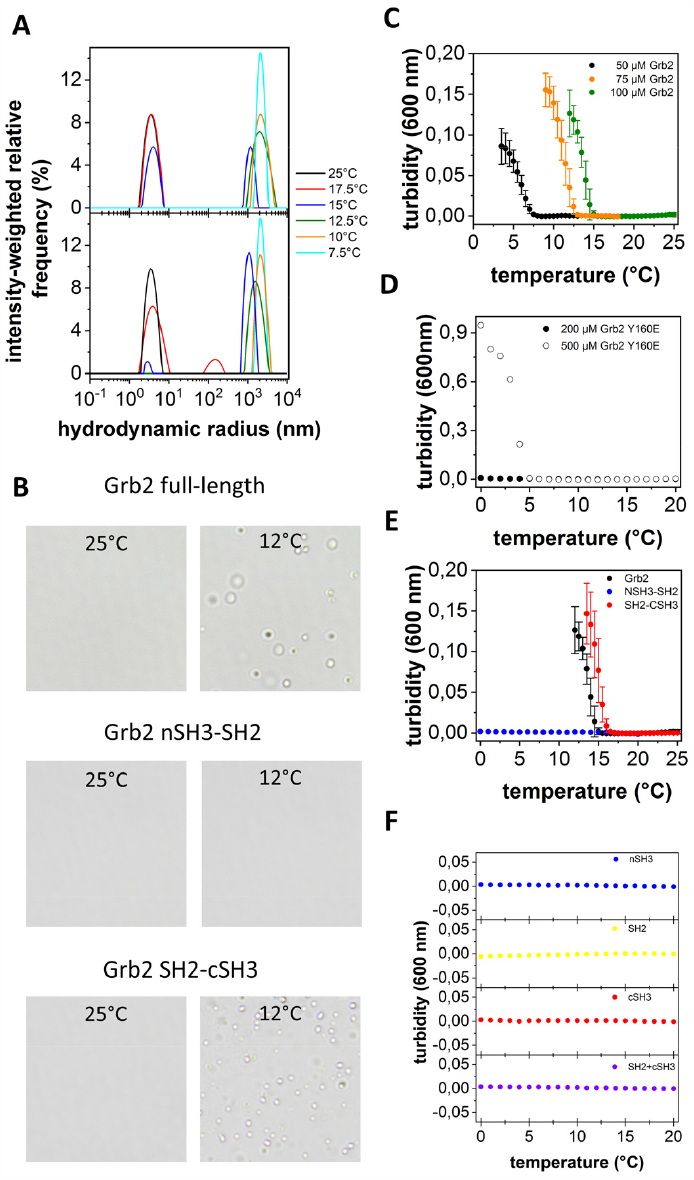
Multimerization of Grb2 in solution. **(A)** Temperature-dependent dynamic light scattering of Grb2. The temperature was lowered stepwise from 25°C to 7.5°C (upper panel) and then increased again to 25°C, revealing reversible assembly of Grb2. **(B)** Polarized light microscopy of Grb2 and the Grb2 di-domains nSH3-SH2 and SH2-cSH3, performed at 25°C and 12°C. Phase separation occurs upon cooling of Grb2 and Grb2 SH2-cSH3. No phase separation is observed for Grb2 nSH3-SH2. **(C-F)** Temperature-dependent turbidity measurements, recorded at different Grb2 concentrations (C), with the Grb2 Tyr160Glu mutant (D), with Grb2 in comparison to the Grb2 di-domains SH2-cSH3 and nSH3-SH2 (E) and with the isolated SH2 and SH3 domains of Grb2 (F).

In the crystal, cSH3 residue Tyr160 is buried in the SH2/cSH3 interface within the Grb2 multimer. Turbidity measurements performed with the pTyr mimic Grb2 Tyr160Glu showed complete suppression of LLPS using the concentration range applied for the wild type (Figure 4D). Increases in turbidity could only be observed at very high Tyr160Glu Grb2 concentrations (500 μM), with a low cloud point temperature below 5°C. To test if the SH2/cSH3 interaction is not only necessary but also sufficient for multimerization, we analyzed separately the tendency of the two Grb2 di-domain constructs nSH3-SH2 and SH2-cSH3 to assemble. The nSH3-SH2 construct did not show LLPS, whereas the SH2-cSH3 di-domain phase separated like full-length Grb2 with a similar cloud point (Fig. 4E). Neither the isolated Grb2 cSH3 and SH2 domains nor a mixture of the two exhibited phase separation, showing that both domains need to be covalently bound for multimerization (Fig. 4F). The turbidity data are therefore in agreement with the Grb2 multimer structure seen in the crystal, demonstrating that an intact SH2/cSH3 interface is needed for Grb2 assembly in solution.

### Phase separation of Grb2-peptide complexes

As the Grb2 multimers are bridged in the crystal by the bidentate Gab1 peptide to form an infinite two-dimensional network, phase separation of Grb2 should be augmented by addition of bridging ligands. Indeed, mixing Grb2 with the bidentate Gab1_472-566_ resulted in spontaneous phase separation at room temperature, without having to decrease the temperature (Fig. 5A). Time courses of turbidity changes as a function of Gab1_472-566_/Grb2 ratio showed that phase separation increased up to a Gab1_472-566_/Grb2 stoichiometry of 0.8:1 and strongly diminished at higher ratios (Figure 5B). This indicates that bridging of Grb2 can occur to a nearly complete 1:1 stoichiometry, despite steric constraints caused by the short linker between the two Grb2 binding motifs on Gab1_472-566_. Further increase of the Gab1_472-566_ concentration showed a decrease in phase separation in line with supersaturation of Grb2 leading to monovalent Gab1 binding.

**Figure 5.**
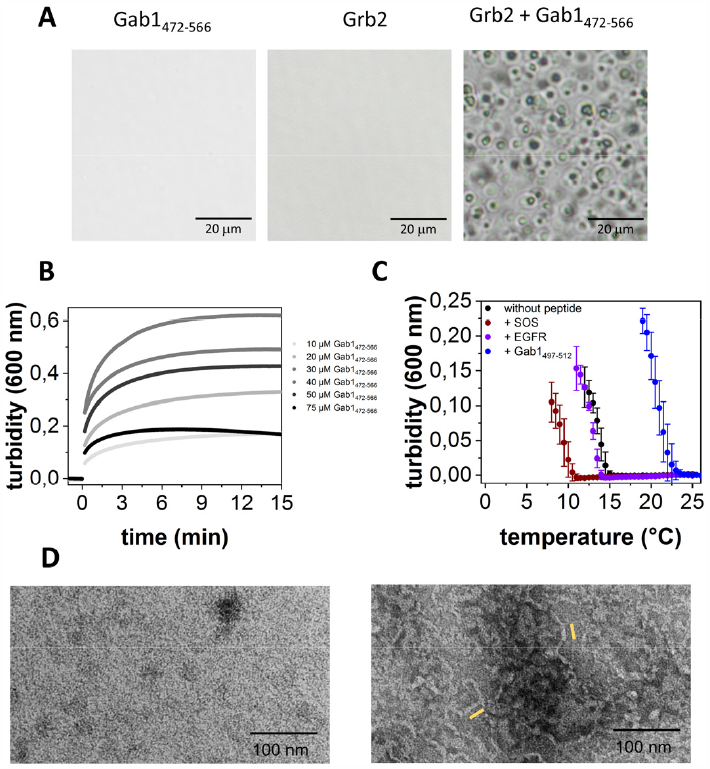
Phase separation of Grb2 – peptide complexes. **(A)** Polarized light microscopy of a Grb2/Gab1_497-528_ mixture in of the single components, performed at 25°C. **(B)** Turbidity measurements of the Grb2-Gab1_497-528_ complex at different Gab1_497-528_ concentrations (concentration of Grb2: 50 μM). **(C)** Temperature-dependent turbidity measurements of Grb2 in presence of stoichiometric amounts of SOS1_1147-1158_, EGFR_1085-1100_ and Gab1_472-566_. **(D)** Transmission electron microscopy of Grb2 (left image) and Grb2 in presence of the nSH3 binding peptide Gab1_497-512_(right image). The width of the yellow bars corresponds to the approximate diameter of the Grb2 filament in the crystal (50Å).

Addition of the Grb2 SH2-binding peptide EGFR_1085-1100_ did not influence phase separation (Fig. 5C), in agreement with the Grb2 SH2 binding site being freely accessible in the Grb2 multimer. Similarly, phase separation still occurred in the presence of Grb2 nSH3-binding peptide SOS1_1147-1158_. Interestingly, the short Gab1_497-512_ peptide containing the Grb2 nSH3 binding motif alone strongly fostered phase separation, shifting the cloud point to > 22°C. Negative stain TEM images of Grb2 incubated at 5°C yielded images of protein dense surfaces, whereas images of Grb2 in the presence of Gab1_497-512_ revealed numerous filament-like structures superimposed on the dense protein lawn (Fig. 5D). The ordered structures exhibited a width comparable to the diameter of the multimeric Grb2 filament in the crystal structure, indicating that Grb2 can form filament-like structures in solution.

## Discussion

The crystal structure determination of the Grb2-Gab1_497-528_ complex sheds light on numerous aspects of peptide recognition by Grb2, a key protein in tyrosine kinase mediated signal transduction. Both consensus (PxxPxR) and non-consensus (RxxK) ligand binding properties of the two Grb2 SH3 domains are utilized, with Gab1 _501_PxxPR_506_ binding to the nSH3 domain in an extended fashion and Gab1 _521_RxxK_524_ forming multiple interactions in the S3 binding site. The Grb2-cSH3 domain structure exhibits high similarity to other Grb2-cSH3 crystal structures (rmsd of superimposed atoms: 0.32-0.52 Å) – including the five residue cSH3 nSrc-loop that adopts the same conformation in all structures and is fixed by three strong hydrogen bonds between nSrc-loop residues and residues of the β-strands B and C as well as one intra-loop hydrogen bond. This stands in sharp contrast to the nSrc-loop structure of the Grb2-nSH3 domain. Whereas the nSrc-loop is well defined and interacts with Gab1 residues R_507_PV_509_ in the Grb2-Gab1_497-528_ complex, it is highly variable in NMR structures of the SOS1_1149-1158_

Grb2-nSH3 complex, disordered in the Grb2 dimer structure and adopts differing crystal packing influenced conformations in structures of ligand-free Grb2-nSH3 (29, 46-48), suggesting inherent flexibility of this loop. The nSrc-loop conformation observed in the present structure resembles that of a ligand-free Grb2-nSH3 structure in which the proline-rich C-terminus of a crystallographic neighbor interacts with the S3/nSrc-loop region, suggesting that this might be a preferred conformation for peptide bound Grb2 nSH3 (Fig. 2E). In addition to the strongly binding SOS1 V_1149_PPPVPPRRR_1158_ sequence, several segments of the SOS1 PR region have been identified that bind to Grb2 nSH3 with dissociation constants in the micromolar range (19, 26). Molecular dynamics simulations of peptides derived from these regions in complex with Grb2-nSH3 indicate that the nSrc-loop varies from an outwardly bent conformation in the ligand-free state to more closed conformations for the simulated peptide complexes (49) resembling the nSH3 nSrc-loop conformation of the Grb2-Gab1_497-528_ complex, which suggests that the nSrc-loop conformation observed in the Grb2-Gab1_497-528_ crystal structure may be a general feature of Grb2-nSH3 ligand binding (Fig.2E). The nSrc-loop structure could also play a role in the elevated phase separation and the induction of filament-like structures that we observed after addition of the monodentate nSH3 binding peptide Gab1_497-512_ to Grb2, with the exact mechanism awaiting further studies. In addition to the Grb2 nSH3 Gab1 interaction, the nSrc-loop is also utilized for extended/exosite binding in other SH3 complex structures (for example in the p47^phox^/p67^phox^:p40^phox^-SH3 or the PEP:Csk-SH3 complexes) (20, 21, 50-52). Binding studies show that dynamin interacts with Grb2-nSH3 via exosite binding including Glu40, positioned at the end of the C-β-strand (22, 23). In view of the Gab1_497-528_ Grb2 structure, this binding mode of dynamin could be realized by binding to an ordered nSrc-loop with interactions including the β-strand C as seen e.g in the p47^phox^:p40^phox^ interaction.

Multivalency is an important driver of phase separation and the basis of a generally accepted concept borrowed from polymer physics that describes LLPS by a network of stickers and spacers (53, 54). Pinpointing these multiple valences to single amino acid residues in protein-driven LLPS, however, has rarely been reported (55, 56). In the present study, we could bridge this gap by the structure determination of the Grb2-Gab1_497-528_ complex and a detailed in-solution analysis of Grb2 and Grb2-peptide condensates. The crystal structure shows the formation of extended multimeric chains formed through cross-linking interactions of the two linear SH3-binding motifs of Gab1_497-512_ with their respective Grb2-SH3 domains. Such short linear motif - domain interactions have been shown to form phase condensates in solution studies, performed with different natural and artificial SH3-domain systems (57). Formation of a two-dimensional mesh as observed in the crystal requires a higher degree of multivalency, however, that we have found to lie in the hitherto unknown ability of Grb2 to self-assemble. Self-assembly is mediated by the intermolecular domain-domain interaction between Grb2-cSH3 and the SH2* domain of a second Grb2 molecule, each of which are covalently linked to their own SH2 and cSH3* domains, respectively, leading to an infinite head-to-tail Grb2 multimer. The same SH2-cSH3* inter-monomeric contacts are observed in the dimer to result in a compact closed dimer structure instead of an infinite filament-like chain as found in our crystals. These results add to the few cases of homotypic multimers involved in phase separation that have been characterized structurally until now, including single-domain head-to-tail interactions of Dix or SAM domains and multimerization of KSHV-dimers by α-helical leucine zipper interactions (58, 59).

The Grb2 homo-multimerization that we detected in the crystal and in solution explains several observations obtained from other studies with phase condensates at supported membranes and / or assemblies using other interaction partners. Clusters of phosphorylated LAT, Grb2 and SOS1, reconstituted at supported lipid bilayers, did not form in the absence of the Grb2-cSH3 domain (37), in line with GRB2-cSH3/SH2 multimers supporting cluster formation. Membraneless cytoplasmic granules that contain fusion oncoproteins of intracellular domain of RTKs like ALK or RETs depend on the presence of Grb2 but not on that of SOS1 and Gab1 (60). After depletion of Grb2, granule formation could be partially rescued by the P49L/P206L Grb2 double mutant that is mutated in both SH3 binding sites. This mutant is unable to bind SH3 ligands yet capable of multimerization as described here, indicating that Grb2 multimers are necessary for granule formation. In addition, our work provides a structural explanation for the formation of pLAT-Grb2 and pEGFR_tail_-Grb2 clusters at supported membranes and of their disruption by phosphorylation of Grb2 Tyr160 as reported by Lin *et. al* (38). In contrast to the Grb2 dimer, multimerization of Grb2 is compatible with SH2-phosphotyrosine binding and therefore the likely oligomeric form of Grb2 in the Grb2-pEGFR_tail_/pLAT condensates at the membrane. Moreover, multimeric Grb2 is able to bind in super-stoichiometric amounts to membrane-bound pEGFR_tail_/pLAT, providing a basis for signal amplification (34).

The Grb2 SH2 and cSH3 domain are connected by an eight-residue linker that is partially disordered in the Grb2-Gab1_497-528_ crystal structure and flexible in the Grb2 monomer in solution (25). Combined with the observation that the Grb2-Grb2 interaction is dominated by the SH2-cSH3 interface in the crystal structure and that multimerization already occurs in the Grb2 di-domain with only SH2 and cSH3 domain present, it is likely that the Grb2 multimer can adopt a more open structure at the membrane that is held together by SH2/cSH3 interactions and retains conformational freedom by the flexible SH2-cSH3 linkers. The change in the assembly behavior of Grb2 observed after addition of the nSH3 binding peptide Gab1_497-512_ suggests that Grb2 multimers with different degrees of compaction exist, ranging from the dense filament-like structure seen in the crystal to an open flexible structure as described above, and that Grb2-multimer density could be regulated by ligand binding. In this way, the Grb2-multimer can offer multiple valences at adaptable positions and with varying density to upstream and downstream interactions partners, making multimerization an important factor in Grb2-mediated signal transduction.

## Materials and Methods

### Expression and purification of Grb2

Human Grb2 and C32S/C198A Grb2 were cloned into a modified pET15b vector with a TEV protease cleavage site. As the cysteine-free Grb2 mutant has been reported to retain essentially the same structure and binding affinities it was used for crystallization and solution experiments in this study. The wildtype Grb2 protein was prepared in parallel and used as control in DLS, turbidity measurements and light microscopy. No significant differences compared to the mutated C32S/C198A Grb2 were observed. In the crystal structure, the two residues are neither located in the ligand-Grb2 or the multimer interface nor do they induce any conformational changes compared to the reported wildtype Grb2 / Grb2-domain structures.

The Grb2 plasmids were transformed into *E*.*coli* BL21(DE3) cells for expression. Protein expressions were carried out in TB medium after induction at an A_600_ of 0.7 with 0.5 mM IPTG over night at 25 °C. After this, bacteria were centrifuged, resuspended in IMAC buffer (50 mM sodium phosphate, 200 mM sodium chlorid, 20 mM imidazole, 1 mM DTT, pH 8.0) with added protease inhibitors, disrupted by sonification and clarified by centrifugation at 48.000 x g at 25 °C. The supernatant was loaded onto a HiTrap IMAC column (Cytiva), washed with IMAC buffer and eluted using a linear gradient with up to 350 mM imidazole. Grb2-containing fractions were pooled and treated with recombinant TEV protease for His_6_-tag cleavage. A second IMAC column was used to separate cleaved and non-cleaved protein. Cleaved protein was pooled and buffer was changed to 20 mM sodium phosphate, pH 6.8, using a HiPrep 26/10 desalting column (Cytiva). The protein solution was then loaded on a HiTrap Q HP column and washed with 20 mM sodium phosphate, pH 6.8. Elution was started with 3 column volumes (CV) of 100 mM NaCl, followed by a linear salt gradient with up to 300 mM NaCl in binding buffer over 15 CV. Finally, all clean fractions of Grb2 were loaded on a size exclusion chromatography column S75 (Cytiva), equilibrated in 20 mM MOPS (pH 7.2), 100 mM NaCl, to remove higher oligomeric species. All monomeric Grb2 fractions were pooled and concentrated with Vivaspin 20 (Sartorius AG). The purification was analyzed by SDS-PAGE and the protein concentration was determined by UV/Vis spectroscopy at 280 nm using a Jasco J-650 spectrometer.

### Expression and purification of single and double Grb2 domains

The Grb2-NSH3_1-57_, Grb2-CSH3_152-217_, Grb2_1-156_ (NSH3-SH2) and Grb2_54-217_ (SH2-CSH3) were cloned from full length Grb2 with flanking primers into pET28d vector with a thrombin cleavage site using the NdeI and BamH1 restriction sites. The plasmids were transformed into *E*.*coli* BL21(DE3) cells. Expressions were carried out in TB medium, or M9 medium for NMR isotope labeled protein. The latter contained ^15^NH_4_Cl and, if necessary, ^13^C-glucose. Protein expression was induced at an A_600_ of 0.7 with 0.5 mM IPTG for 3 h at 37 °C. Cells were resuspend in IMAC buffer, lysed by sonication and purified from soluble material. The supernatant was loaded on a HiTrap IMAC column (Cytiva) and washed with IMAC buffer, before the buffer was changed to 20 mM Tris/HCl, 150 mM NaCl, pH 8.0. Thrombin was then added to the column for His_6_-tag cleavage overnight. Afterwards, the column was washed again with IMAC buffer to elute all cleaved products. The two single domain proteins were further purified using a HiLoad Superdex 75 size-exclusion chromatography column (Cytiva). The purified domains were dialyzed against NH_4_HCO_3_ and lyophilized. The two double domain variants were subjected to a buffer change into 20 mM sodium phosphate, pH 7.5 for Grb2_1-156_ and pH 8.0 for Grb2_54-217_ using a HiPrep 26/10 desalting column (Cytiva) after the second IMAC step and subsequently further purified using a HiTrap Q HP column. For this, protein was loaded and elution was started with 3 CV of 100 mM NaCl, followed by a linear gradient of up to 300 mM NaCl in binding buffer over 15 CV. Finally, all clean fractions of Grb2 were also loaded on a size exclusion chromatography column S75 (Cytiva) equilibrated in 20 mM MOPS, 100 mM NaCl pH 7.2 to remove higher oligomeric species.

### Crystallisation and structure determination

Grb2 was concentrated to 13 mg/mL and mixed with Gab1_497-528_ peptide in a ratio of 1:1.5. Crystallisation was performed at 13°C by hanging drop vapor diffusion by adding 1 μL of protein-peptide solution to 1 μL of 0.1M Tris/HCl (pH 8.0), 22% PEG4000 and 10% 2-propanole. Monoclinic crystals with dimensions of 100 × 60 × 30 μm^3^ appeared after 14 days. Diffraction data was collected at 100K on BESSY beamline 14.2 (Helmholtz Zentrum, Berlin, Germany) using a Pilatus 2M detector (Dectris). Diffraction data were integrated and scaled using XDS. Initial phases were determined by molecular replacement with the program PHASER using high resolution structures of the separate Grb2-cSH3 and SH2 domains (PDB codes: 2vwf and 3wa4) as well as the coordinates of Grb2-nSH3 of the full-length ligand-free Grb2 structure (1gri) as search models (18, 29, 61). Model building and structure refinement were carried out using the programs COOT and PHENIX, respectively. Structure visualization was performed with PyMOL (PyMOL Molecular Graphics System, Schrödinger) and interaction analysis with PISA.

### Isothermal titration calorimetry (ITC)

ITC titration curves were recorded with a ITC200 calorimeter (GE Healthcare) at 25°C. All measurements were done in 20 mM MOPS (pH 7.2), 100 mM NaCl, unless stated otherwise. To avoid buffer dilution effects, all samples were extensively dialyzed. 50 μM of Grb2 were used and 500 – 800 μM of Gab1–peptides unless otherwise stated. Protein concentrations were determined by UV absorption. ITC stirring speed was set to 700 rpm and the feedback mode was set to “high”. The ITC was equilibrated long enough to ensure that no major baseline drift occured before the measurements. For data analyses, the manufactoring software (Origin 7 SR4 v7.0552) was used. Resulting peaks were baseline-corrected as needed and integrated to acquire the quantity of heat generated. A single binding model to obtain the *K*_*D*_-values and stoichiometries was used to fit the titration curve.

### In vitro liquid - liquid phase separation assay

Temperature dependent turbidity assays were conducted using a UV-Vis spectrometer equipped with a Peltier temperature controller. The protein concentration was 100 μM, dissolved in buffer with 20 mM MOPS (pH7.2), 100 mM NaCl. All samples were first equilibrated at 37 °C to ensure non-multimeric protein and then transferred into a quartz curvet with a 1 cm path length. Samples were then cooled down in 0.5 °C steps and equilibrated for 5 min at every step to reach equilibrium. Thereafter, the absorptions at A_600_ were measured as an indicator for turbidity. When phase separation occurs, the sample changes from clear to turbid when cooled down below the phase transition temperature. For DLS measurements a Litesizer 500 (Anton Paar) was used. Samples are irradiated with a 40 mW semiconductor laser at 658 nm and detection at a 90° angle. Grb2 protein concentration was 100 μM in 20 mM MOPS (pH 7.2), 100 mM NaCl. The protein was directly filtered into quartz cuvettes (Hellma Analytics) and equilibrated for 5 min at each temperature. The resulting correlation function was analysed using the Kalliope Software. Imaging of temperature-dependent phase transitions was performed with a polarisation microscope BX10 (Olympus) in transmission mode. Sample conditions and treatments are identical to the turbidity assays. The measurements were done in self-prepared glass chambers.

### NMR analyses

NMR experiments were performed on a Bruker Avance III 800 MHz spectrometer equipped with a CP-TCI cryoprobe at 25 °C. Samples contained 1 mM nSH3 domain in 20 mM sodium phosphate (pH 7.2) with 10% (v/v) D_2_O and sodium trimethylsilylpropanesulfonate (DSS) used for chemical shift referencing. For backbone assignment standard triple resonance experiments (HNCA, HNCACB, HN(CO)CACB and HNCO) were used. Gab1_497-541_ or SOS peptide were added to an excess of 1:1.2. The complex formation was monitored by 2D ^1^H-^15^N HSQC spectra with a WATERGATE sequence for water suppression. The chemical shift differences were calculated according to:

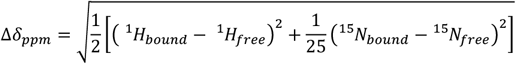

### Negative-stain transmission electron microscopy

EM grids were prepared at 5°C by loading 10 μL of the protein suspension (0.05 mg/mL) onto glow-discharged copper TEM grids with continuous 10–12 nm carbon film coating (300 mesh size; Quantifoil Micro Tools, Großlöbichau, Germany). Excess liquid was blotted off with a strip of filter paper after 45 s followed by staining with 10 μL 2% (w/v) aqueous uranyl acetate solution. Specimens were dried and examined in an EM 900 transmission electron microscope (Carl Zeiss Microscopy, Oberkochen, Germany), and micrographs were recorded with an SM-1k-120 slow-scan charge-coupled device (slow-scan CCD) camera (TRS, Moorenweis, Germany).

## Abbreviations used

SH2: Src Homology 2 domain
SH3: Src Homology 3 domain
PH: Pleckstrin homology domain
PR region: proline rich region
IDR: intrinsically disordered region
rmsd: root mean square deviation
RTK: receptor tyrosine kinase
LLPS: liquid liquid phase separation

## Acknowledgements

This research was supported by grants BA1821/6-1 (JB) and FE439/7-1 (SF) as well as the RTG 2467 “Intrinsically Disordered Proteins – Molecular Principles, Cellular Functions, and Diseases” (project number 391498659, JB, SF and MTS) of the Deutsche Forschungsgemeinschaft (DFG). ML and TG were funded in part by the EU/ESF program HAL-OX (Halle – Oxford Research Network “Disease Biology and Molecular Medicine”, project no. ZS/2016/08/80642). KM was supported by the Medical Faculty of the Martin Luther University Halle-Wittenberg and the Heads Up Cancer Foundation (Oxford, GB). MTS and JB acknowledge funding from the Federal Ministry for Education and Research Germany BMBF (project 03Z22HN22), and JB the European Regional Development Funds EFRE for significant investments into the NMR infrastructure. SF would like to thank the Structural Genomics Consortium (Oxford, GB) for helpful discussions. CB and MTS thank the Helmholtz-Zentrum Berlin for the allocation of synchrotron radiation beam time and would like to thank the staff at beamline 14.2 at the BESSYII storage ring for excellent support.

