## Supplemental Figure 1 for "Self-assembly of Grb2 meshworks revealed by Grb2-Gab1_497-528_ complex structure"

**Figure S1**

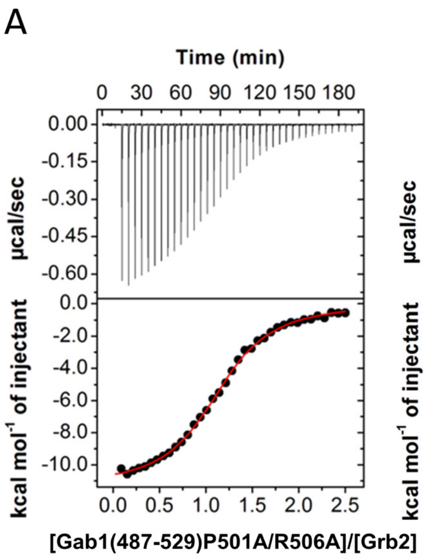


***Figure S1***

***ITC measurements of Gab1-peptides titrated to Grb2 and Grb2 domains.*** **(A)** Titration of the peptide P501A/R506A Gab1$_{487\text{-}529}$, in which the PxxPxR motif has been mutated to Grb2 **(B)** Titration of the bidentate peptide Gab1$_{497\text{-}541}$ to Grb2 cSH3. **(C)** Titration of SOS variant peptide PPPLPPRRRR to Grb2 nSH3 (62). The ITC data that is in line with a single binding mode and the resulting *Kd* value are in agreement with published values, confirming the structural integrity of the binding site of the purified Grb2 nSH3 domain.
